# C1q/MASP complexes – hybrid complexes of classical and lectin pathway proteins are found in the circulation

**DOI:** 10.1101/2024.03.15.584944

**Authors:** Anne Rosbjerg, Tereza Alica Plchová, Rafael Bayarri-Olmos, Bettina Eide Holm, Ida Sandau Pedersen, Mikkel-Ole Skjoedt, Peter Garred

**Affiliations:** Department of Clinical Immunology, Laboratory of Molecular Medicine, Copenhagen University Hospital, Copenhagen, Denmark; Institute of Immunology and Microbiology, University of Copenhagen, Copenhagen, Denmark

**Keywords:** Complement system, MASP-2, MASP-3, C1q, factor D, alternative pathway

## Abstract

Complement pathways, traditionally regarded as separate entities in vitro, are increasingly noted for cross-communication and bypass mechanisms. Among these, the MBL/Ficolin/CL associated serine protease-3 (MASP-3) — a component of lectin pathway pattern recognition receptors (PRMs) — has shown the ability to process critical substrates like pro-factor D and insulin growth factor binding protein-5 (IGFBP-5). Given shared features between lectin pathway PRMs and C1q from the classical pathway, we hypothesized that C1q might be a viable in vivo binding partner for the MASPs.

We used microscale thermophoresis, ELISA and immunoprecipitation assays to detect C1q/MASP complexes and functionally assessed the complexes through enzymatic cleavage assays.

C1q/MASP-3 complexes were detected in human serum and correlated well with MASP-3 serum levels in healthy individuals. The binding affinity between MASP-3 and C1q in vitro was in the nanomolar range, and the interaction was calcium-dependent, as demonstrated by their dissociation in the presence of EDTA. Furthermore, most of the circulating C1q-bound MASP-3 was activated. Based on immunoprecipitatin, also C1q/MASP-2 complexes appereared to be present in serum. Finally, C1q/MASP-2 and C1q/MASP-3 in vitro complexes were able to cleave C4 and pro-factor D, respectively.

Our study reveals the existence of C1q/MASP complexes in the circulation of healthy individuals and both C1q/MASP-2 and C1q/MASP-3 complexes display proteolytic activity. Hence, this study uncovers a crosstalk route between complement pathways not previously described.

## Introduction

The complement system is the immune system’s vanguard, poised to detect and confront foreign invaders or malfunctioning host cells. This highly sophisticated proteolytic cascade acts to eliminate undesired entities and restore tissue balance. At the core of this system are proteins known as pattern recognition molecules (PRMs), which act as the sensory apparatus of the immune system. They detect foreign, unfamiliar structures on cellular membranes, effectively acting as a trigger for the domino effect of the complement cascade through associated enzymes (Ricklin et al. 2010).

The complement system functions in a complex network of protein interactions, set in motion by three key pathways:the classical, lectin, and alternative pathways. Each pathway is characterized by unique complexes. In the classical pathway, the activation complex is C1, which includes the PRM C1q and serine proteases C1r and C1s. Meanwhile, the lectin pathway has a selection of PRMs:Three ficolins (ficolin-1,-2 and-3) and three collectins (mannose-binding lectin (MBL), CL-10 and CL-11). They sit in complexes with MBL/ficolin/CL-associated serine proteases (MASPs) (Garred et al. 2016). The MASPs are produced from two genes, *MASP1* and *MASP2*, which undergo alternative splicing to generate five different gene products. From *MASP1*, two enzymes are generated, MASP-1 and MASP-3, along with a third truncated protein called MAP-1 (also known as MAp44), which lacks serine protease activity. *MASP2*, on the other hand, produces one enzyme known as MASP-2 and another truncated protein without protease activity called MAP-2 (also known as MAp19). The alternative pathway functions as an amplification loop, which depends on the rate-limiting protein factor D (FD) (Garred et al. 2016; Ricklin et al. 2010).

The classical and lectin pathways bear a striking resemblance in their architecture. The PRMs are constructed by trimers of collagen-like stalks and binding domains that assemble into higher oligomers (Teillet et al. 2005; Thielens et al. 2017). This enables the molecules to have multiple binding sites towards their targets. The complexes between lectin and classical pathway PRMs and their respective serine proteases are assembled in a very similar manner with the heavy chains of the serine proteases facing inwards and interacting with the collagen-like stalks in the PRM molecule and their light chains pointing outwards. In these complexes, the MASP and C1r/C1s heavy chains form dimers and tetramers, respectively (Chen and Wallis 2001; Kishore and Reid 2000). The binding sites of the MASPs and C1r/C1s are likely conserved, and the binding properties are, therefore, highly comparable (Phillips et al. 2009).

The similarity between pathways continues as the lectin and classical pathway have identical substrates; C4 and C2 are cleaved by both C1s and MASP-2, and MASP-1 cleaves C2 (Kishore and Reid 2000; Matsushita et al. 2000). Cleavage of C4 and C2 results in the formation of the C3 convertase C4b2b (previously known as C4b2a) and from this point, the pathways converge (Bohlson et al. 2019).

Interactions and crosstalk between complement pathways and other systems, such as coagulation, are well documented, gradually unraveling a complex network of associations. Emerging evidence suggests that these boundaries are not as rigid as previously believed. An intriguing example is the interplay between the lectin and alternative pathways, with MASP-3 from the lectin pathway showing an essential role in the alternative pathway through its cleavage of pro-FD (Dobó et al. 2016; Hayashi et al. 2019). Additionally, MASP-3 is proteolytically active against insulin growth factor protein 5 (IGFBP-5), implying its relevance to numerous biological processes (Cortesio and Jiang 2006).

Considering the remarkable similarities and close interactions between the classical and lectin pathways, we propose a novel hypothesis:classical and lectin pathway molecules may’mix and match’ during complex formation. With this intriguing possibility in mind, this study explores whether MASPs can form complexes with C1q – a potential revelation for our understanding of immune system dynamics.

## Results

### In vitro complex formation between C1q and MASPs

We initially established whether C1q and MASPs could form complexes in vitro using purified and recombinant proteins. In an ELISA system with immobilized immune complexes and C1q, C1q was able to bind the following proteins:A) rMAP-1, B) rMASP-1, C) rMASP-3 and D) a mix of purified MASP-1,-3 and MAP-1. The complexes assembled in a concentration-dependent manner, as shown in fig. 1.

**Figure 1.**
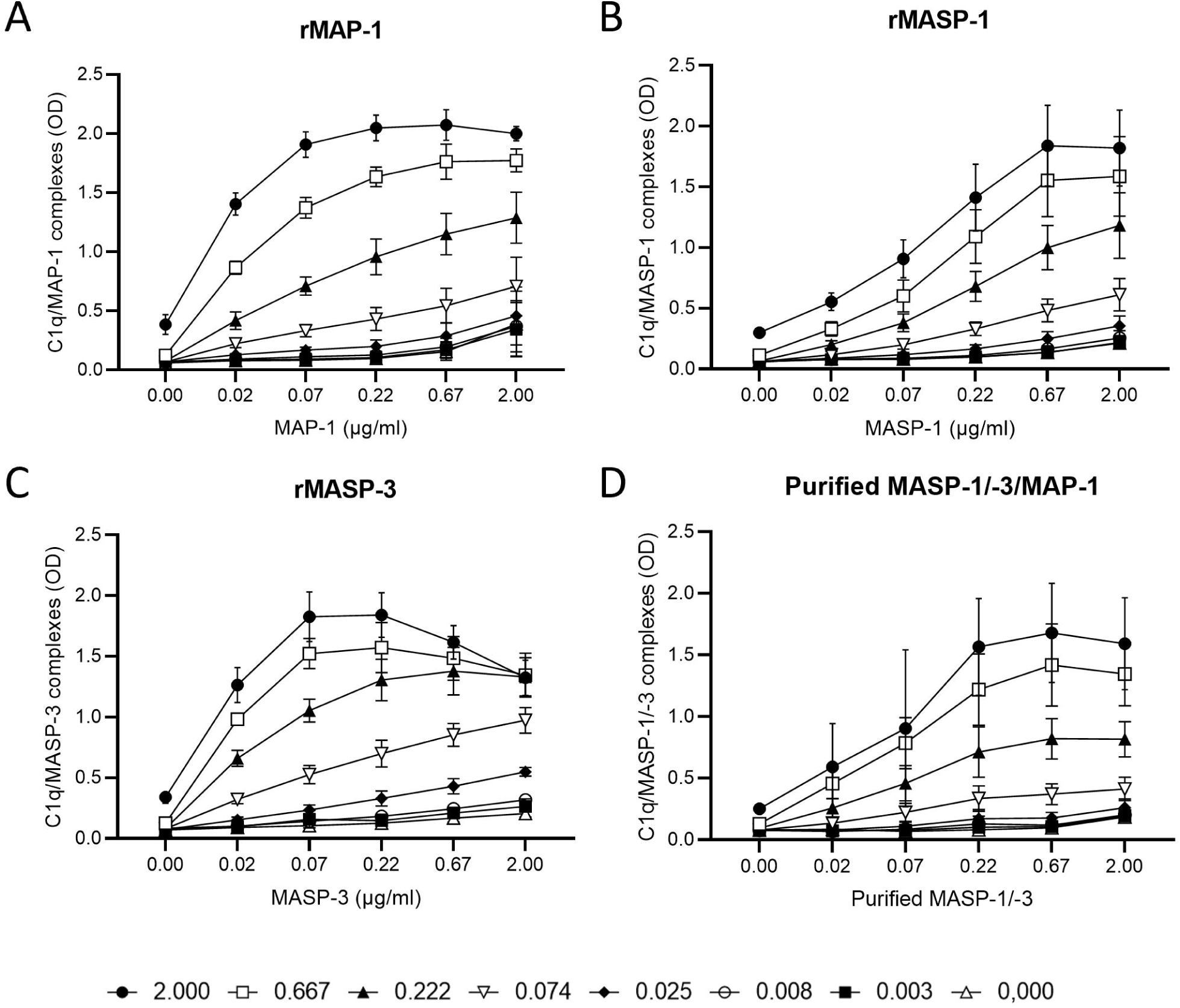
In vitro C1q/MASP complexes. Formation of C1q/MASP complexes were performed in ELISA on human albumin/anti-human albumin immune complexes. A titration of purified C1q was added to the immune complexes first and afterwards a titration of recombinant or purified MASP was added. A mAb against the common heavy chain of the MASPs (8B3) was used as detection. A) C1q/rMAP-1, B) C1q/rMASP-1, C) C1q/rMASP-3 and D) C1q/pur. MASP-1/-3/MAP-1. The curves show a C1q and MASP concentration dependent assembly of the proteins. The results represent the means of three independent experiments and the error bars show the standard deviation.

### Binding affinity between C1q and MASP-3

Next, we investigated the biophysics of the in vitro complexes. Using Microscale Thermophoresis (MST), we measured the binding affinity between C1q and MASP-3 in order to compare it with the known MASP-3 ligand MBL. Fluorescently labelled recombinant MASP-3 was mixed with either purified C1q or rMBL and the binding affinities of fluid phase complexes were assessed. The Kd values of MBL/MASP-3 were 1.79 nM (1.22-2.65 nM), whereas C1q/MASP-3 had a Kd value of 69.56 nM (56.27-85.98 nM) (fig. 2).

**Figure 2.**
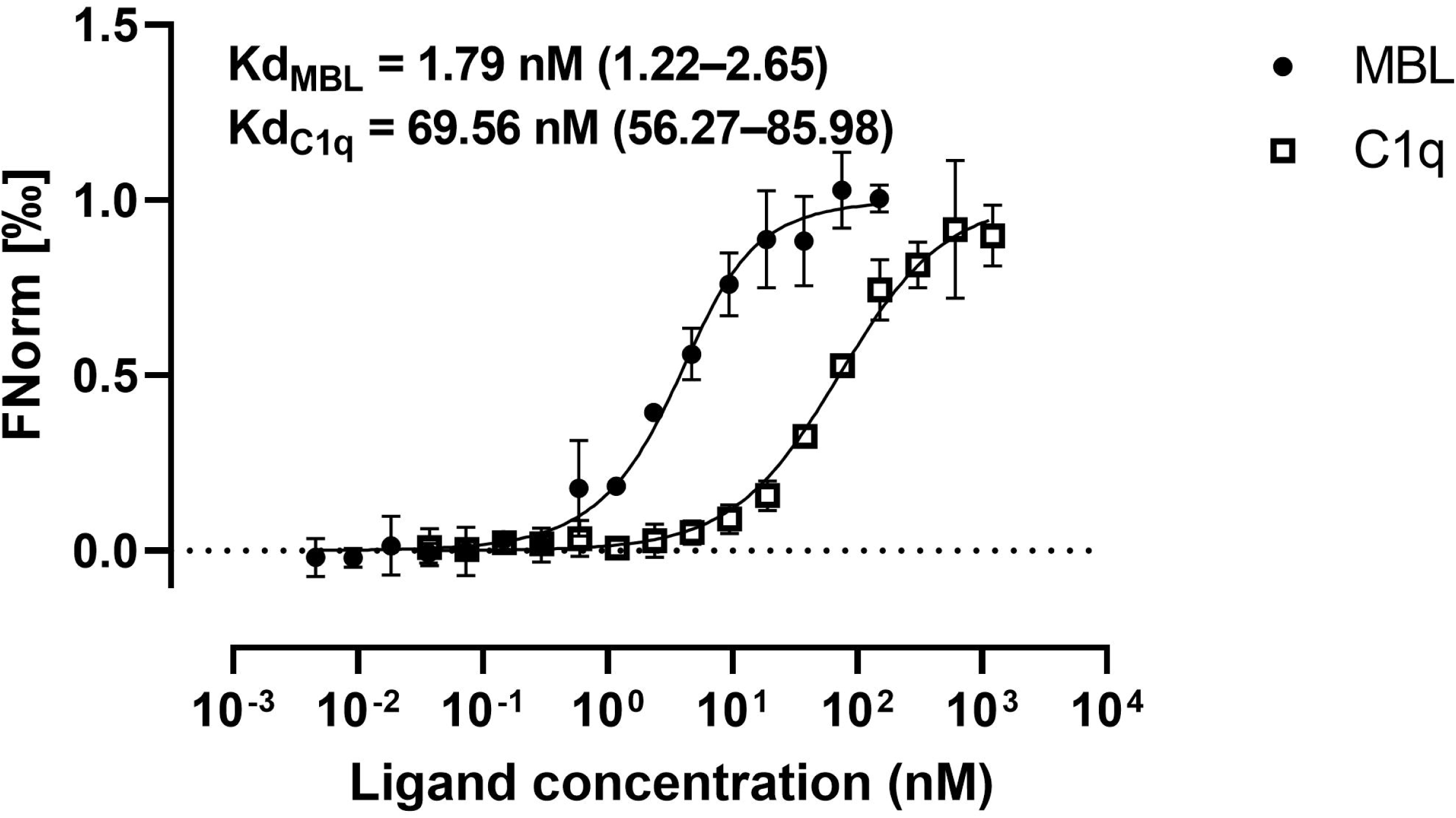
MST dose response curves for rMASP-3 binding to rMBL and C1q. MST traces from three independent experiments were used to calculate binding affinities (Kd) between rMASP-3 dimers and rMBL or pur. C1q. Data are presented as fraction bound.

### C1q/MASP-3 serum complexes

The existence of endogenous C1q/MASP-3 complexes was investigated in a sandwich ELISA with the principle of catching MASP-3 and detecting C1q. To avoid direct C1q binding to the coated anti-MASP-3 antibody (which would cause a false positive result), we used F(ab’)2 fragments of the coated antibody. Using this setup, C1q/MASP-3 complexes were detected in normal human serum (NHS) (fig. 3a). EDTA disrupted the binding, indicating that these complexes were calcium-dependent (fig. 3a). To make sure we were not detecting C1q being directly bound the coated antibody, we tested the binding of a high concentration of C1q in a buffer with either Ca^2+^ or EDTA. Binding of C1q did not differ between buffers with or without EDTA (suppl. fig. 1), hence, the EDTA mediated loss of signal was a result of C1q/MASP-3 complex disruption and not a false positive detection of C1q. We reached the same conclusion when coating with F(ab’)2 fragments from an antibody recognizing the common heavy chain of MAP-1, MASP-1 and MASP-3 (fig. 3b). Hence, C1q/MASP-1 complexes may also form. We observed no binding to F(ab’)2 fragments from an irrelevant mouse antibody (fig. 3c) neither to non-coated plates (fig. 3d).

**Figure 3.**
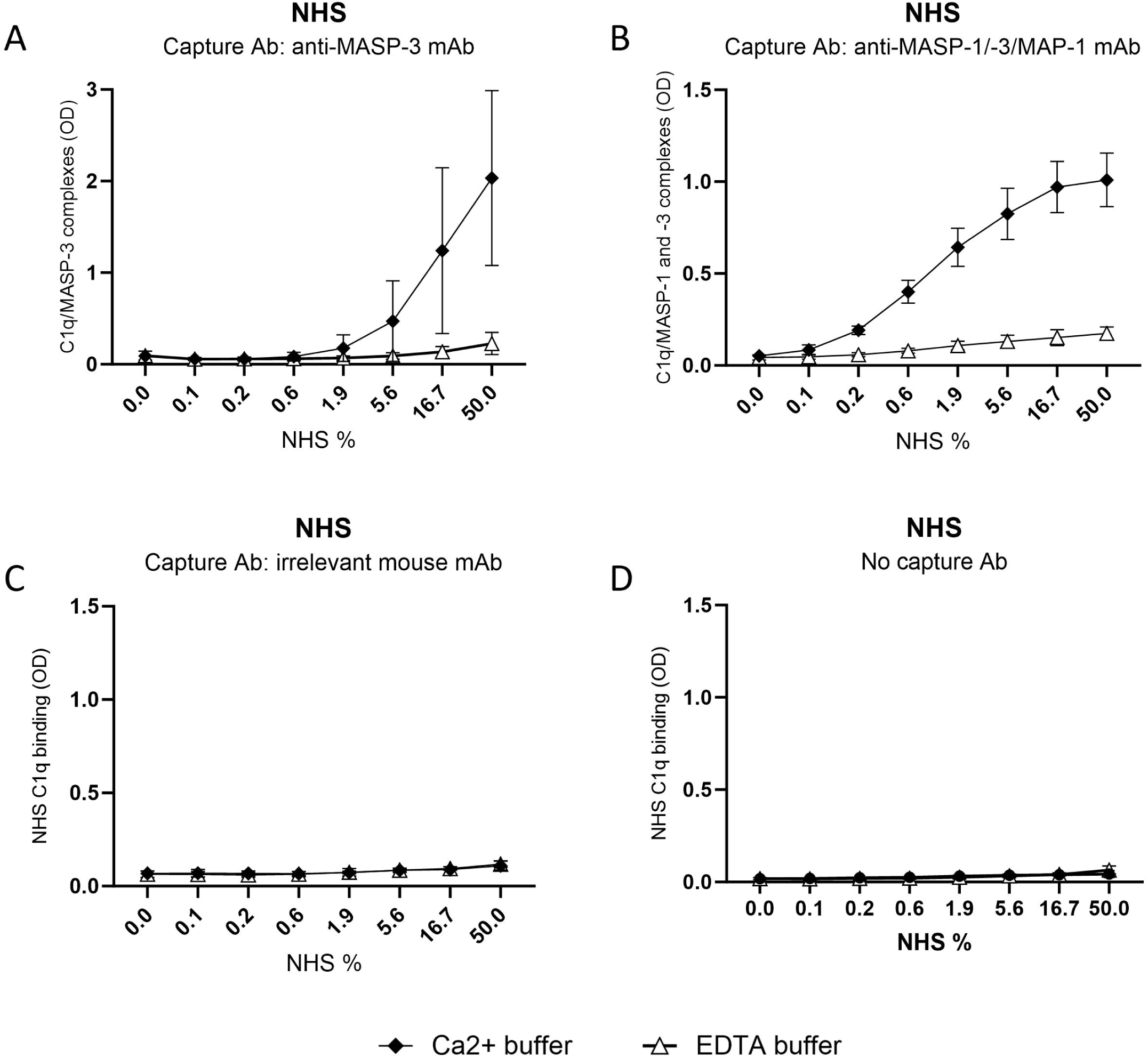
C1q/MASP *in vivo* complexes. C1q/MASP complexes were measured in a human serum pool with/without pre-incubation with 10 mM EDTA. The assay is a sandwich ELISA assay coating with anti-MASP mAb F(ab’)2 fragments and detecting with an anti-C1q pAb. The difference between A-C is the coating Ab; A) anti-MASP-3 mAb (7D8) F(ab’)2, B) anti-MASP-1/-3/MAP-1 mAb (8B3) F(ab’)2, C) Irrelevant mouse mAb F(ab’)2 and D) No coating Ab. Graphs A and B show detection of calcium dependent complexes on the coated anti-MASP mAbs, which is not seen on the control mouse mAb (C). The results represent the means of three independent experiments and the error bars show the standard deviation.

### Immunoprecipitation of C1q/MASP serum complexes

Immunoprecipitation of C1q/MASP serum complexes was performed with streptavidin-coupled magnetic beads incubated first with biotinylated F(ab’)2 from either an anti-MASP-2 mAb, anti-MASP-3 mAb or an anti-MASP-1/-3/MAP-1 mAb as bait for MASP complexes. After incubation in serum, the beads were eluted, and a western blot confirmed the presence of C1q in the eluate of all anti-MASP mAb F(ab’)2 fragment pull-downs and not the negative control (fig. 4a top panel). Hence, C1q forms complexes with the MASPs in circulation under physiological conditions. Detection of MBL was included in a parallel experiment shown in the bottom panel as a validation of the assay. Interestingly, MBL/MASP-3 complexes did not appear to present in this context. The low concentration of MBL in serum is likely the reason that MBL is not detected.

**Figure 4.**
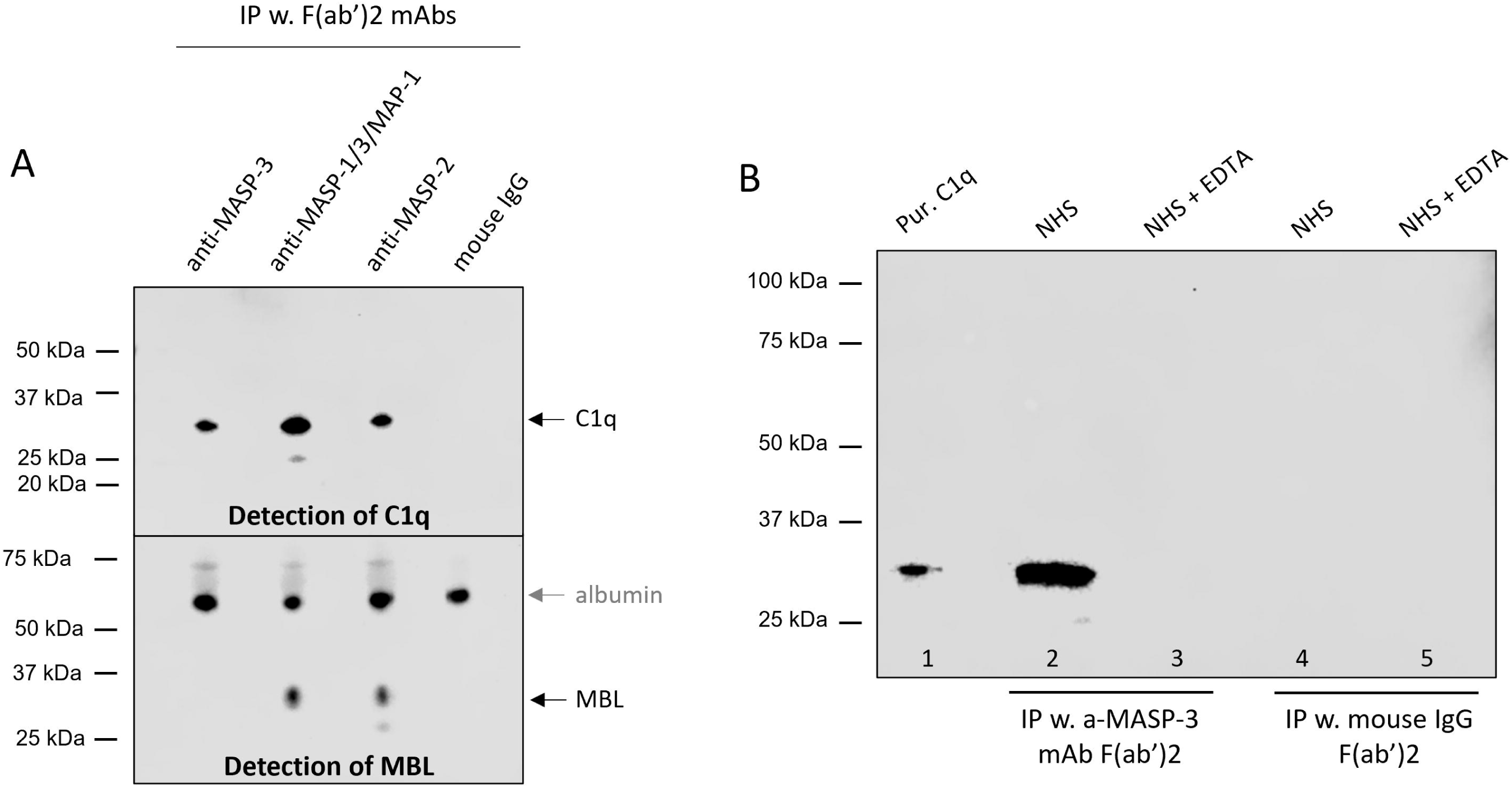
Immunoprecipitation of *in vivo* C1q/MASP complexes. A) C1q/MASP-2 complexes were immunoprecipiated using biotinylated anti-MASP mAb F(ab’)2 and streptavidin-coupled magnetic beads. The following mAb F(ab’)2 fragments were used for pull down:Lane 1) anti-MASP-3 mAb (7D8) F(ab’)2. Lane 2) anti-MASP-1/-3/MAP-1 mAb (8B3) F(ab’)2. Lane 3) anti-MASP-2 mAb F3-5 F(ab’)2. Lane 4) Irrelevant mouse mAb F(ab’)2. The eluate from the beads were analyzed by western blotting detecting C1q (top panel) and MBL (bottom panel). B) C1q/MASP-3 complexes were pulled down on an ELISA plate coated with an anti-MASP-3 mAb F(ab’)2 or an irrelevant mouse mAb F(ab’)2. The plates were eluted well to well and the accumulated eluate was applied in SDS-PAGE/western blotting to detect C1q. Lane 1) Purified C1q, lane 2) IP of NHS with the anti-MASP-3 mAb (7D8) F(ab’)2, lane 3) IP of NHS/EDTA with the anti-MASP-3 mAb (7D8) F(ab’)2, lane, 4) IP of NHS with control mouse mAb F(ab’)2, lane 5) IP of NHS/EDTA with control mouse mAb F(ab’)2. The C1q detection in lane 2 shows that C1q/MASP-3 complexes were immunoprecipitated by the anti-MASP-3 mAb F(ab’)2 and the absence of C1q in lane 3 shows the calcium dependency. The blot is a representative of two independent experiments.

We next confirmed the presence of C1q/MASP-3 in serum with a plate-based immunoprecipitation assay. NHS was incubated on a plate coated with anti-MASP-3 mAb F(ab’)2 fragments and the bound serum proteins were afterward eluted and analyzed by western blotting. As shown in fig. 4b, we detected C1q in the elution fraction (lane 2), meaning that a part of the immobilized serum MASP-3 was in complex with C1q. It was also confirmed that the complexes were assembled in a calcium-dependent manner, as the C1q band was missing under calcium chelating conditions (lane 3). Again, F(ab’)2 fragments from an irrelevant mouse IgG was used as a negative control (lane 4-5). In general the plate-based assays did not work well with the anti-MASP-2 mAb F(ab’)2 fragments.

### MASP-3 in a C1 complex preparation

We then investigated whether C1 complexes purified from serum would contain MASP-3. Hence, 1 and 2 µg of C1 (Comptech) was loaded on an SDS gel and subsequently blotted to a membrane in order to detect MASP-3 with a specific anti-MASP-3 mAb. Zymogen rMASP-3 was loaded as a positive control and C1q (Comptech) as a negative control. Indeed, the C1 preparation contained MASP-3, as visualized in fig. 5a in lane 2 and 3. C1q in lane 4 did not generate a MASP-3 band. Hence, MASP-3 is only present when the full C1 complex is purified. In fig. 5b the samples were run under reducing conditions, which caused the MASP-3 bands in lane 2 and 3 to almost vanish. This indicates that MASP-3, in complex with C1q, is in its activated state where heavy and light chain are only held together by a disulfide bond and thus split under reducing conditions. High amount of light chain is necessary to get a visible band with this antibody; hence, this may explain why we do not see a band at ∼35 kDa.

**Figure 5.**
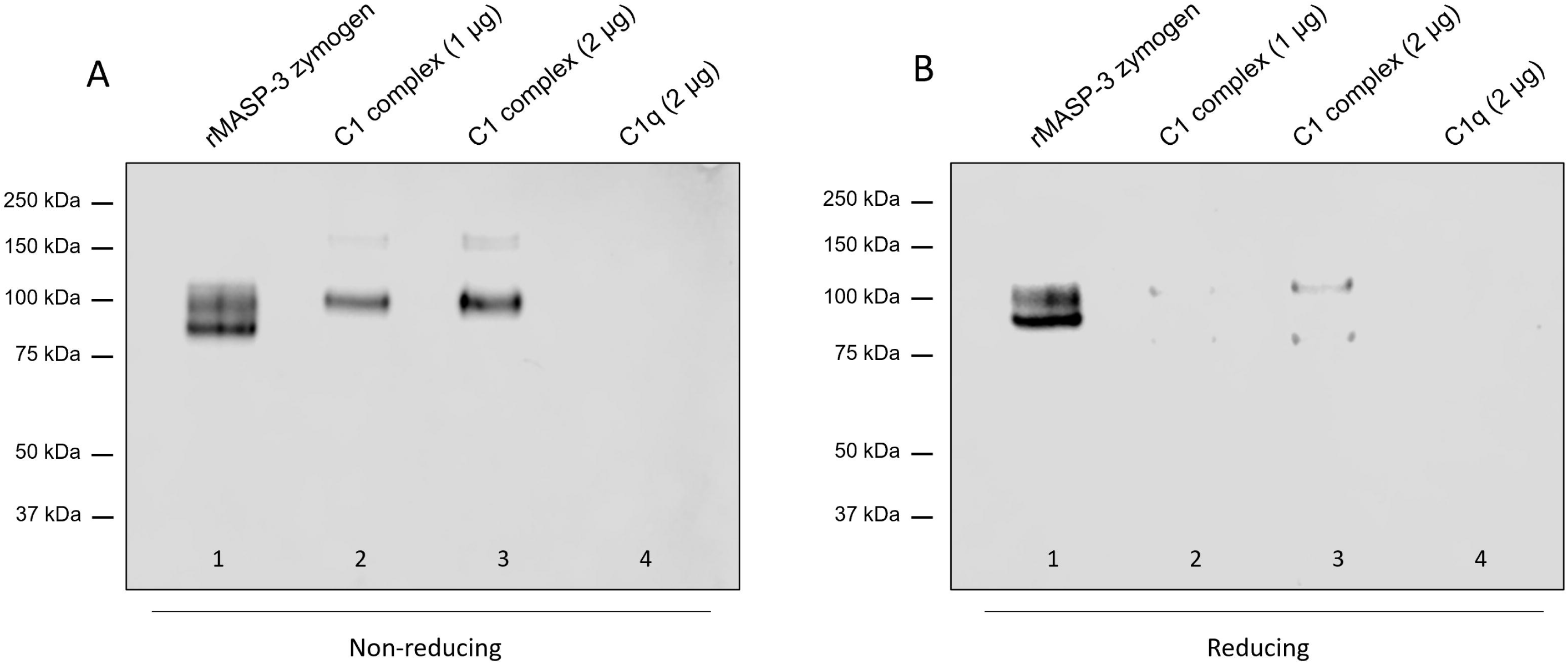
MASP-3 in purified C1 complexes. The presence of MASP-3 in a preparation of purified C1 complexes were analyzed by western blotting using an anti-MASP-3 mAb (38:12-3) for detection. The gel was run under A) non-reducing and B) reducing conditions. Lane 1) rMASP-3, lane 2) 1 µg purified C1, lane 3) 2 µg purified C1 and lane 4) 2 µg purified C1q.

### C1q/MASP-3 complex levels in a healthy cohort

After having established that C1q/MASP-3 complexes are present in the circulation, we investigated the occurrence on a population level. Using the sandwich ELISA to detect serum complexes, we measured the level of complexes in a cohort of 99 healthy donors (fig. 6a). C1q and MASP-3 were also measured separately (fig. 6b-c). The level of C1q/MASP-3 complexes are presented in arbitrary units on a log scale, and it shows that there is a wide distribution on the number of complexes in the cohort. Moreover, the levels of C1q/MASP-3 complexes were correlated with MASP-3 (fig. 6d) and not C1q (fig. 6e).

**Figure 6.**
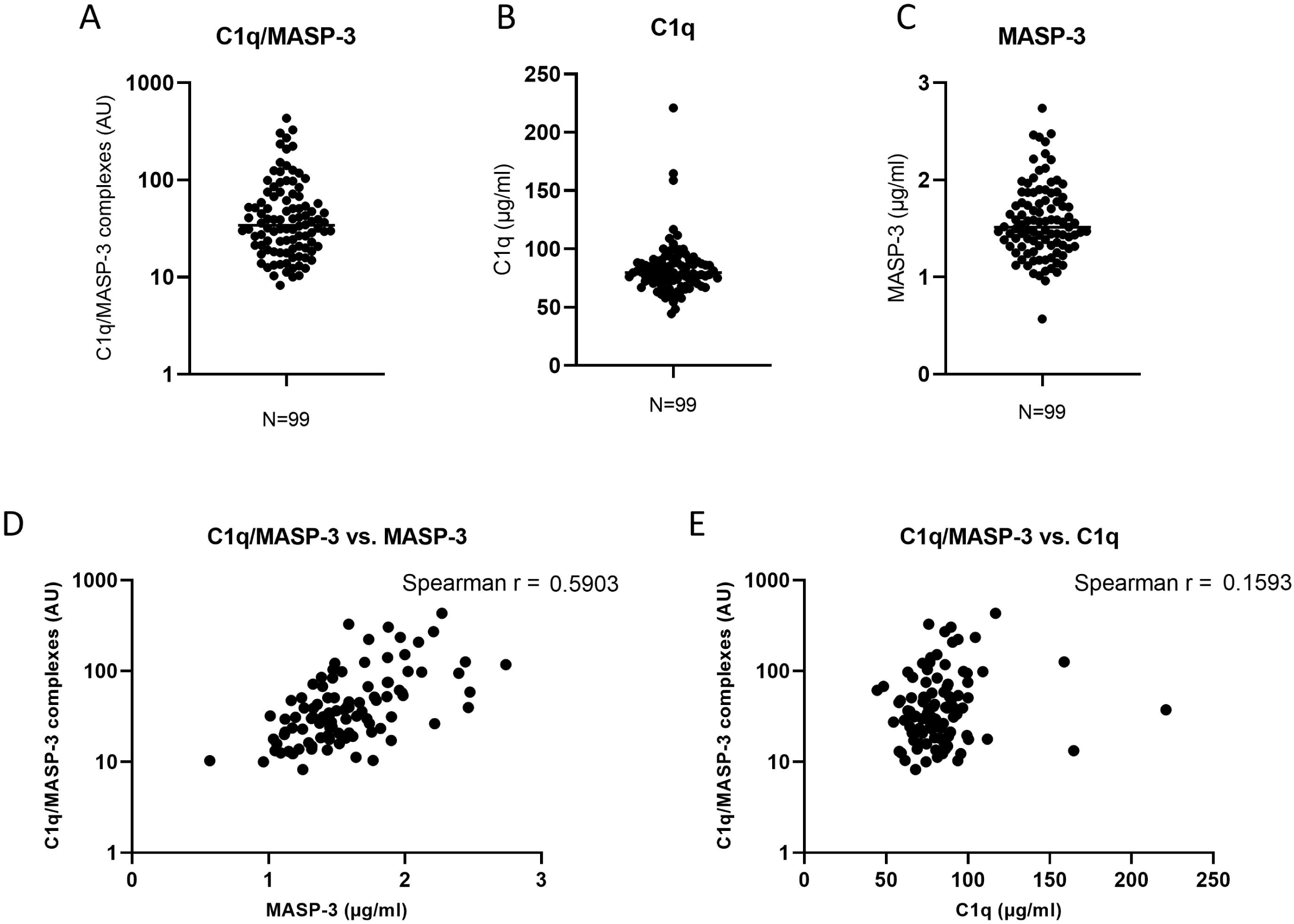
C1q/MASP-3 complexes in a cohort of healthy donors. The content of C1q/MASP-3, C1q and MASP-3 were measured in serum samples of 99 healthy donors. A) C1q/MASP-3, B) C1q, C) MASP-3, D) correlation between C1q/MASP-3 and MASP-3 and E) correlation between C1q/MASP-3 and C1q.

### C1q/MASP-3 cleavage of FD

The enzymatic functionality of C1q/MASP-3 complexes was tested in a pro-FD cleavage assay. C1q/MASP-3 complexes bound to immune complexes in ELISA were incubated with recombinantly expressed pro-FD at different time intervals. Afterwards, the supernatant was analyzed in a western blot to check the activation status of FD, which was possible with an in-house mAb recognizing only mature FD. As seen in the top panel in fig. 7, pro-FD was cleaved into mature FD over time by C1q/active MASP-3 complexes, whereas C1q/zymogen MASP-3 only showed a minor cleavage similar to the buffer control. Total FD was also detected as shown in the bottom panel. Suppl. fig. 2 shows the specificity of the mAb recognizing only mature FD, which was made recombinantly in CHO cells. Pro-FD was produced in the presence of protease inhibitors in the culture media and was not recognized by the mAb. A mAb against both pro and mature FD demonstrated the overall presence of FD in in the supernatant (suppl. fig 2).

**Figure 7.**
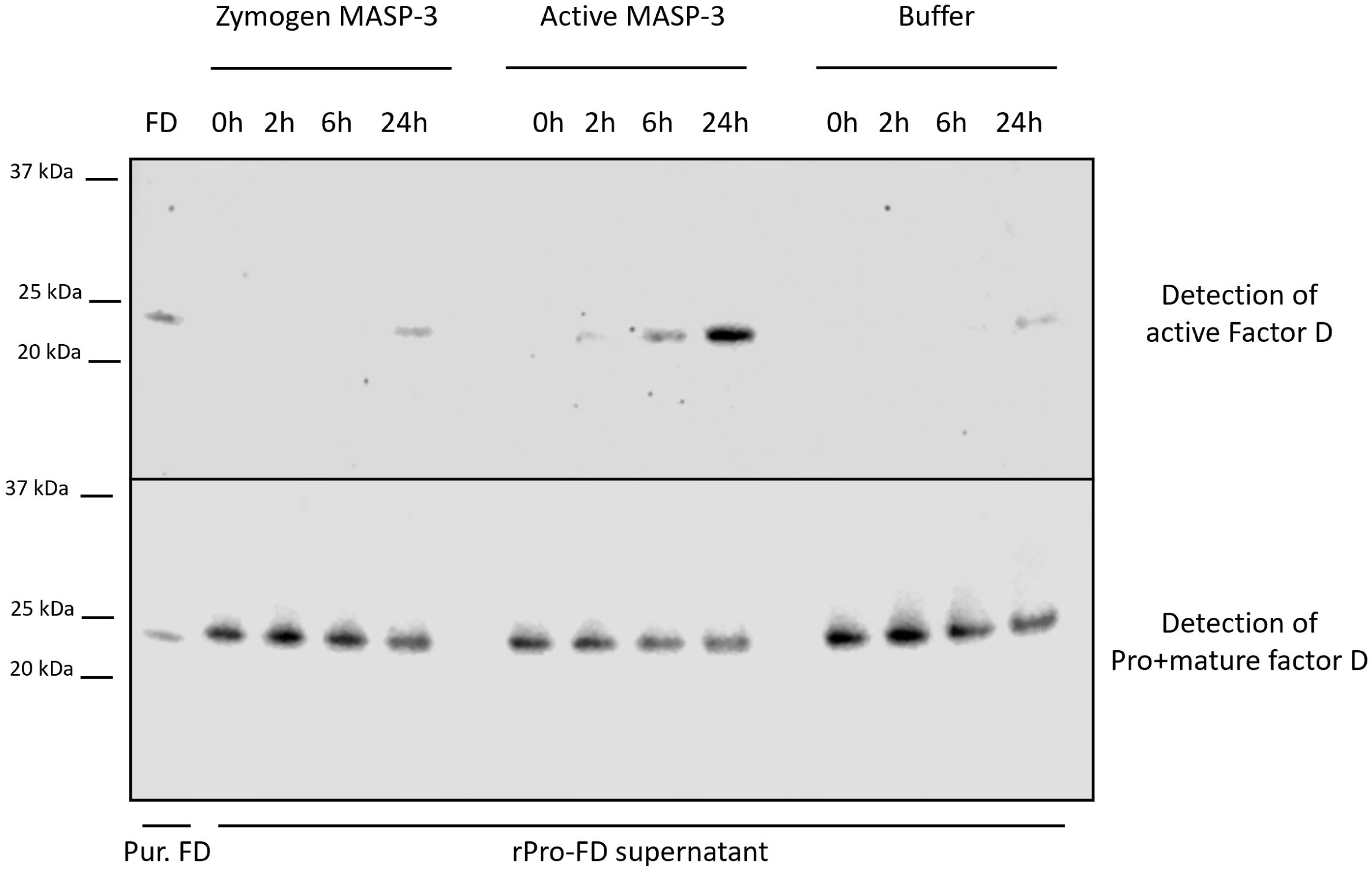
Pro-FD cleavage by C1q/rMASP-3 complexes. Enzymatic functionality of active MASP-3 sitting in complex with C1q was tested in its ability to cleave pro-FD. Pro-FD was added to C1q/rMASP-3 complexes bound to immune complexes in an ELISA plate. The supernatant was recovered after 0, 2, 6 and 24 h and tested in a western blot using an anti-FD mAb (Act 17) only recognizing the cleaved form of FD and not pro-FD (top panel) or a common anti-FD mAb (F1-11) (bottom panel). Complexes with active rMASP-3, zymogen rMASP-3 and only C1q (buffer) were compared. The complexes with active rMASP-3 appeared to be enzymatically active since mature-FD specific bands appeared after 6 and 24 h, whereas the zymogen rMASP-3 was comparable to the buffer control.

### C1q/MASP-2 cleavage of C4

We also tested the functionality of C1q/MASP-2 complexes with a C4 cleavage assay. C1q was first added to an ELISA plate with coated immune complexes followed by rMASP-2 to form C1q/MASP-2 complexes. C4 cleavage were assessed by measuring C4b deposition after adding either purified C4 or C1q depleted serum (fig. 8a). C4b is deposited in the presence of C1q/rMASP-2 and C4, but not when omitting rMASP-2. Moreover, C4b was deposited in a C1q dose-dependent manner. The tendency was the same when C1q depleted serum was used as a C4 source, especially in the lower C1q concentrations (fig. 8b).

**Figure 8.**
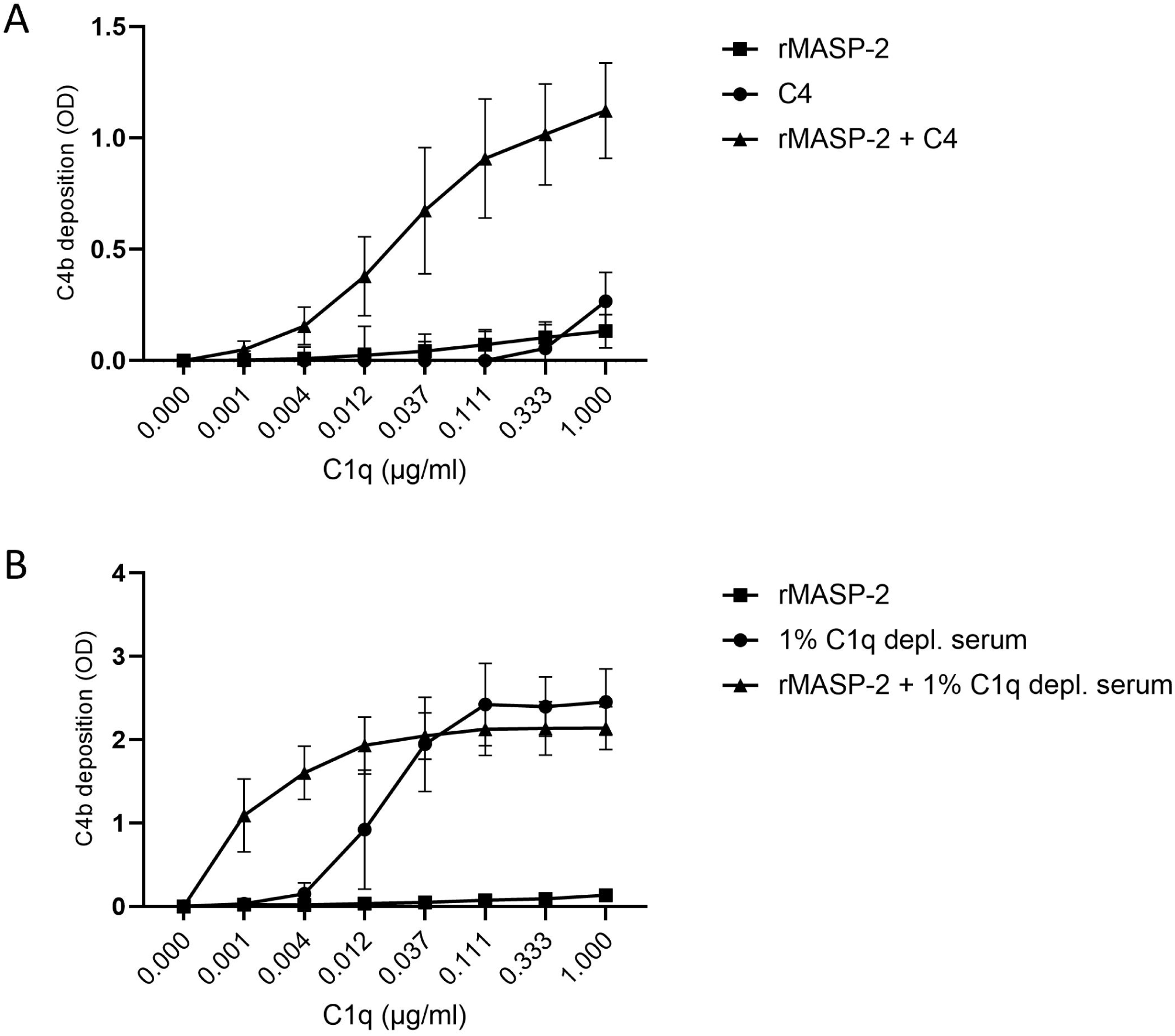
C4 cleavage by C1q/rMASP-2 complexes. C1q/rMASP-2-driven C4 cleavage was assessed in an ELISA assay where rMASP-2 was first allowed to bind to C1q sitting on coated immune complexes before adding either purified C4 or C1q deficient serum. A) C4b deposition via C1q/rMASP-2 complexes using purified C4. B) C4b deposition via C1q/rMASP-2 complexes using C1q depleted serum. The results represent the means of three independent experiments and the error bars show the standard deviation.

### Zymogen-MASP-3 inhibition of classical pathway

Finally, zymogen rMASP-3 was shown to compete with C1r/C1s for C1q binding in a classical pathway ELISA assay, thus hindering complement activation (fig. 9); C4b deposition was hampered when zymogen rMASP-3 was allowed to pre-bind to C1q prior to adding C1q depleted serum. The level of inhibition depended on the concentration of rMASP-3.

**Figure 9.**
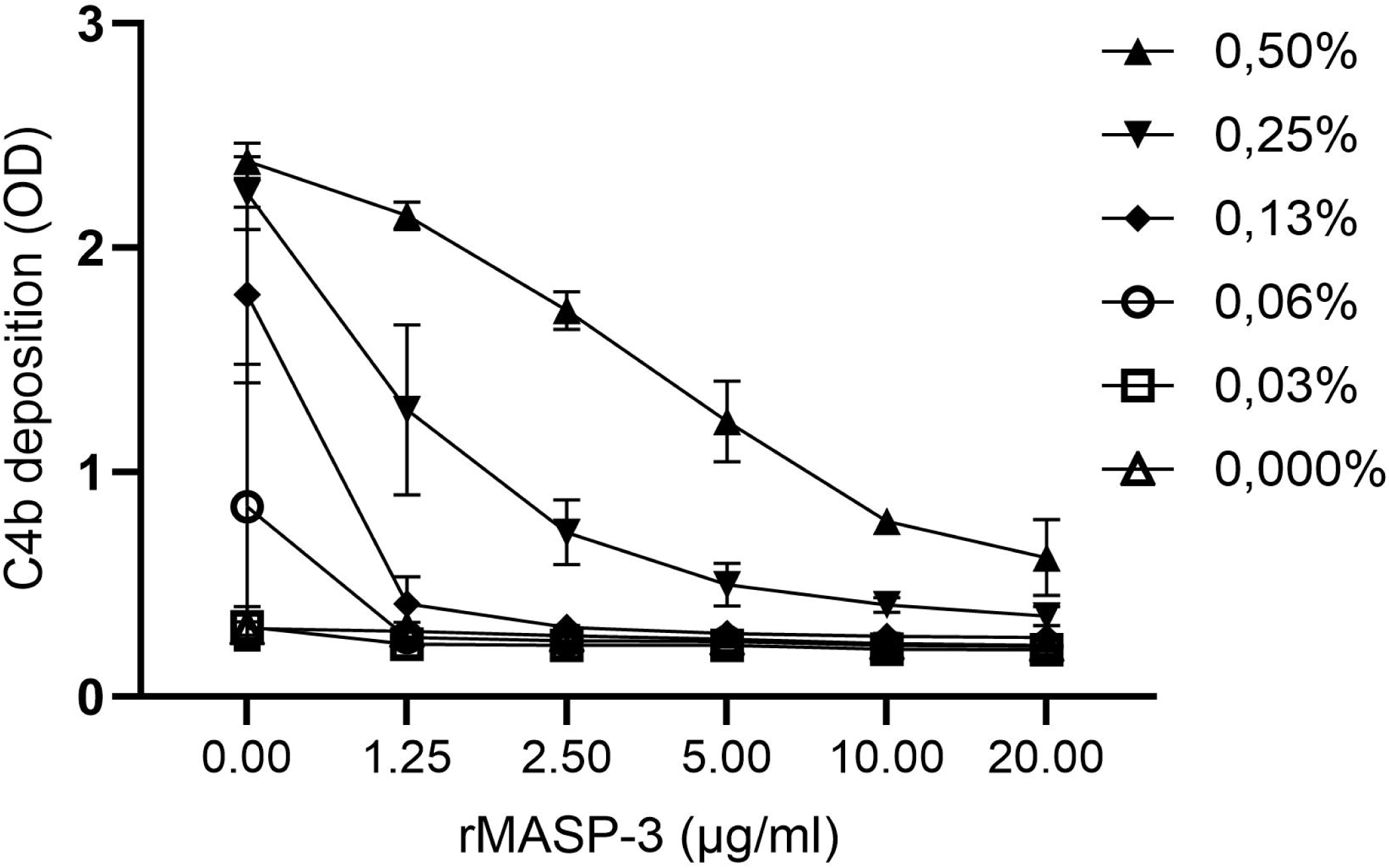
Zymogenic rMASP-3 inhibition of classical pathway. Zymogenic rMASP-3 competition with C1r/C1s for C1q binding was assessed in a classical pathway activation assay, where zymogen rMASP-3 was first allowed to bind to C1q sitting on coated immune complexes before adding C1q deficient serum. The inhibitory effect of zymogen rMASP-3 was shown as a decrease in C4b deposition. The results represent the means of three independent experiments and the error bars show the standard deviation.

## Discussion

With this study, we have discovered a new group of complement protein complexes in human plasma that weaves together the lectin, classical and alternative pathway, namely C1q/MASP complexes.

We have focused mainly on the C1q/MASP-3 complexes. Based on our cohort of healthy individuals, it seems there is a large distribution of C1q/MASP-3 complex levels in the population, mostly dependent on the MASP-3 plasma levels. Others have previously investigated classical/lectin pathway hybrid complexes in vitro, but to our knowledge, this is the first time that C1q/MASP complexes have been shown to happen in vivo (Lu et al. 1990; Ohta et al. 1990; Phillips et al. 2009).

A few studies have previously shown that functional complexes between MBL and C1r/C1s can assemble in vitro using purified proteins (Lu et al. 1990; Ohta et al. 1990) and other studies have shown the opposite with C1q and the MASPs (Phillips et al. 2009). Thiel et al. examined the classical and lectin pathway complexes in human serum; however, they did not detect any hybrid complexes (Thiel et al. 2000). The technical difference between that previous study and our current study is that we pulled down the MASPs and then measure C1q, whereas they did it the other way around and pulled down C1q. In that case, the C1q/MASP complexes are being outnumbered by the C1q/C1r/C1s complexes in the pull-down material. As seen on the western blot in fig. 5, a high concentration of C1 is needed to detect MASP-3, so possibly, the MASPs have been below the detection level in the Thiel et al. study.

The affinity of MASP-3 towards C1q and MBL are both in the nanomolar range, with MBL being the strongest binding partner. The Kd values found in this study closely resemble previously described affinities between MASP-1 and C1q and MBL (Phillips et al. 2009). MASP-1 and MASP-3 have identical heavy chains, meaning that they also have identical binding domains towards the PRMs. Hence, the affinities are comparable. The concentration of MASP-3 dimers in plasma is around 25 nM (Skjoedt, Palarasah, et al. 2010) and thus below the measured Kd value of C1q/MASP-3 complexes (∼70 nM). However, the plasma concentration likely does not represent the local concentration in tissues where complexes might form.

Analogous to the C1 complex and (MBL/ficolin/collectin-11)/MASP complexes, the C1q/MASP complexes were shown to be calcium-dependent. Moreover, we could see that zymogen rMASP-3 pre-bound to C1q competed with C1r/C1s and inhibited C4 cleavage (fig. 9). All in all, this implies that the C1r, C1s and the MASPs are assembling with C1q in the same fashion.

The functionality of the C1q/MASP-3 complexes was demonstrated using pro-FD, which is a known substrate of MASP-3 (Dobó et al. 2016; Oroszlán et al. 2015). For this we used a newly developed mAb recognizing only the cleaved mature factor D and we used recombinant pro-factor D from Flp-In™ CHO cells cultured in the presence of a protease inhibitor cocktail, which prevented the enzymatic cleavage of pro-factor D into mature FD in the culture supernatant. When pro-factor D was added on top of immobilized C1q/MASP-3 complexes, active MASP-3 cleaved pro-FD into FD. Notably, the cleavage kinetics seemed relatively slow (6-24 h). In a study by Dobo et al. from 2016, it is shown that the half-life of pro-FD in serum ex vivo is around 2 h, and they show that inhibition of MASP-3 drastically prolongs the half-life (Dobó et al. 2016). In our study, the protease inhibitors were still present in the pro-FD preparation during the cleavage assays and that might affect MASP-3 activity. In the future, it would be optimal to have a pro-FD preparation free of enzymatic inhibitors. Yet, to our knowledge, this is the first time full-length human rMASP-3 has been shown to cleave pro-FD, as previous studies have mainly worked with the catalytic domain of MASP-3 (Dobó et al. 2016; Oroszlán et al. 2015).

C1q/MASP-2 complexes were also shown to be functional in an in vitro setting – here using a C4 cleavage ELISA assay. The plate-based assay measuring C1q/MASP complexes did not work well with the F(ab’)2 fragments of the anti-MASP-2 mAb. However, we were able to detect the complexes with immunoprecipitation followed by western blotting (fig. 4b), suggesting that all MASPs may form complexes with C1q to a certain extent.

Recent discoveries have placed MASP-3 in a central position in the alternative pathway as a main facilitator of pro-FD cleavage into mature FD (Dobó et al. 2016). However, the framework around this function is still unclear – e.g does the cleavage happen in the fluid phase or via immobilized MASP-3 complexes? In this regard and based on our study, it can be speculated whether C1q, in some cases, works as a scaffold for MASP-3 to activate the alternative pathway under speffic conditions. C1q may also help retain MASP-3 in the circulation as a recent study has suggested that MASP-3 binding to lectin pathway PRMs slows down the clearance of MASP-3 (Kusakari et al. 2022).

With this study, we have established a unique intersection between complement pathways. We conclude that C1q/MASP complexes are found in the circulation and that both C1q/MASP-2 and C1q/MASP-3 complexes are enzymatically functional. With this newly discovered layer of the immune system, the complexity becomes evident, and future studies will determine the physiological impact of C1q/MASP complexes.

## Methods

### Recombinant and Serum Purified Proteins

Recombinant MASPs and MBL were expressed and purified essentially as previously described (Larsen et al. 2004; Skjoedt, Palarasah, et al. 2010). In short, rMBL were expressed in CHO-DG44 cells cultivated in RPMI 1640 medium (Sigma-Aldrich) supplemented with 10% FCS, 100 U/ml penicillin+0.1 mg/ml streptomycin (Gibco, 15-140-122), 2 mM L-glutamine (Gibco, 25030-024), and 200 nM methotrexate. rMASPs were expressed in CHO-DG44 cells cultivated in Power CHO serum-free media (Lonza, Vallensbaek, Denmark) supplemented with 20 U/mL penicillin, 20 μg/mL streptomycin, and 2 mM L-glutamine and 200 nM methotrexate. rPro-FD was produced in Flp-In^TM^-CHO cells (Invitrogen, USA) cultured in Ham’s F12 media containing 10% heat-inactivated FCS (56 °C for 30 minutes), 100 U/ml penicillin, 0,1 mg/ml streptomycin, 0.5 mg/ml Hygromycin B (Invitrogen, 10687010) and ReadyShield Protease Inhibitor Cocktail (PIC) (Sigma-Aldrich, PIC0006). Collected supernatant was used unpurified. rProprotein convertase subtilisin/kexin type 5 (PCSK5) was expressed in the ExpiCHO expression system produced by Gibco™. rPCSK5 was produced with a 17 aa tag from MAP-1 for detection purposes using an anti-MAP-1 mAb 20C4 (Skjoedt, Hummelshoj, et al. 2010). After transfection, ExpiCHO cells were cultured for 12 days according to the manufacturer’s Max Titer protocol (32 °C, 5% CO_2_). Collected supernatant was concentrated 2-fold by centrifugal filtration with a 50 kDa filter (Amicon, UFC805096). MASPs were purified from a normal human serum pool of 3 healthy donors by indirect affinity chromatography using anti-ficolin-3 mAb FCN334 followed by elution with 10 mM EDTA. Purified C1 (A098) and C1q (A099) were purchased from Comp Tech (Texas, USA).

### MASP-3 activation and purification

Zymogen rMASP-3 containing a His-Tag was incubated at 50 µg/ml for 24 h in the up-concentrated ExpiCHO PCSK5 culture supernatant at room temperature (RT) while gently shaking. rMASP-3 was afterwards purified using Ni-NTA resin, dialyzed in PBS overnight (ON), and up-concentrated by centrifugal filtration with a 50 kDa filter to approximately 150 µg/ml.

### Generation of monoclonal Factor D antibody

NMRI mice were immunized subcutaneously three times with 15 µg of the peptide ILGGREA-C OR ILGGREAEAHARPYMA-C coupled onto diptheria toxid and mixed with Gerbu adjuvant P in a 1:1 ratio. Four days before the fusion the mice received an intravenous injection with 15 µg antigen without adjuvant. The SP2/0-AG14 myeloma cell line was used as fusion partner to generate hybridomas and PEG was used as fusogen. Positive clones were selected for reactivity to ILGGREA-C and ILGGREAEAHARPYMA-C and contra-screened against PPRGRILGGREAEAHARPYMA-C all. The antibody was purified from culture supernatant by protein G affinity chromatography using the ÄKTA Pure system with a HiTrap Protein G column (GE Healthcare).

### ELISA measurements of in vitro complexes

The formation of complexes between C1q and MASP-1, MASP-3 and MAP-1 was measured in an ELISA assay where 10 µg/ml recombinant human albumin (rHA) (Sigma-Aldrich, A9731) was coated on Nunc MaxiSorp flat-bottom 96-well plates (Thermo Fisher Scientific, 442404) at 4 °C ON and afterward incubated with rabbit anti-HA antibody 1:2000 (Dako, A0001) 2 h at RT. Then purified C1q (CompTech, A099) was added in a serial dilution starting at 1 µg/ml at 4 °C ON. Recombinant (Skjoedt, Palarasah, et al. 2010) or purified MASPs (Rosbjerg et al. 2014) were added in a serial dilution starting at 2 µg/ml for 1 hr at RT. The binding of MASPs to C1q was measured using a biotin-tagged monoclonal mouse antibody 8B3 (Skjoedt, Palarasah, et al. 2010) that recognizes a common epitope on MASP-1,-3 and MAP-1 followed by HRP-conjugated streptavidin (Merck, RPN1231). The plates were revealed using TMB One as the substrate (Kementec, 4380A). All dilutions and washing were performed in barbital/tween buffer (4 mM sodium barbital, 145 mM NaCl, 2.6 mM CaCl_2_, 2.1 mM MgCl_2,_ 0.05% tween 20, pH 7.4).

### Affinity measurements by microscale thermophoresis

The affinity of MASP-3 towards MBL and C1q in solution was determined by MicroScale Thermophoresis (MST) using a Monolith NT.115 Pico RED (Nanotemper Technologies GmbH, Munich, Germany). rMASP-3 (in-house) was labelled using the Monolith His-Tag Labeling Kit RED-tris-NTA 2nd Generation kit (Nano Temper MO-L018) in barbital/tween buffer. Labelled rMASP-3, C1q (CompTech, A099), and rMBL (in-house) were centrifuged at 17,000 x g for 30 min 4 °C before performing the binding experiments. In the meantime, low-binding dilution plates were blocked with assay buffer (barbital/tween) for 30 min at RT. Two-fold serial dilutions of the target C1q or rMBL (starting concentration 1,220 nM and 150 nM, respectively) were incubated with 4 nM of rMASP-3 in low-binding dilution plates and allowed to interact with gentle shaking in the dark for 30 min at RT. Samples were loaded into Monolith capillaries, and measurements were performed using the Pico mode, high MST-power at LED setting of 25%. Dose response curves from three independent experiments were evaluated using the MO. Affinity Analysis 3 Software (Nanotemper) using a Kd fit model and constraining the target concentration.

### ELISA measurements of serum complexes

The sandwich ELISA assay was performed with 4 µg/ml coated F(ab’)2 fragments that were generated from full-size antibodies using the Pierce Mouse IgG1 Fab and F(ab’)2 Preparation Kit (Thermo Fisher, 44980) according to the manufacturer instructions. To measure C1q/MASP-1 and-3 complexes, we used F(ab’)2 of the anti-MASP-1/-3/MAP-1 mAb 8B3 and to specifically measure C1q/MASP-3 complexes we used F(ab’)2 of an anti-MASP-3 mAb 7D8 (Skjoedt, Palarasah, et al. 2010). Before adding a normal human serum (NHS) pool (10 donors) to the plates, the NHS pool were pre-treated with 10 mM EDTA in PBS or only PBS ON at 4 °C. The NHS and NHS/EDTA was added in serial dilutions starting in 50% and purified C1q was added in a serial dilution starting at 25 µg/ml. NHS was diluted in TBS/T/Ca (Tris-HCL, 150 mM NaCl, 2.5 mM Ca^2+^, 0,05% tween 20, pH 7,4) and NHS/EDTA was diluted in TBS/T/EDTA (Tris-HCL, 150 mM NaCl, 10 mM EDTA, 0,05% tween 20, pH 7,4). C1q was diluted on both buffers. NHS and C1q were incubated on the plates 1H at RT. Afterwards, a biotin-tagged polyclonal anti-C1q Ab (Agilent, A0136) was incubated for 1 hr at RT followed by HRP-coupled streptavidin 1:10000 and TMB One.

### Immunoprecipitation on streptavidin-coupled magnetic beads

A total of 5 µg F(ab’)2 fragments generated from anti-MASP-3 mAb 7D8, anti-MASP-1/-3/MAP-1 mAb 8B3, anti-MASP-2 mAb F3-5 (Peter et al. 2022) and an irrelevant mouse IgG1 mAb (BD Biosciences, 557273) were coupled to to Dynabeads M-280 streptavidin (Thermo Fisher, 11205D) by incubating 30 min at RT. The beads were washed 5 times in PBS/0.1% BSA and then incubated with 1ml NHS (no clot activator) mixed with 1 ml Tris-buffered saline (TBS) with Ca^2+^ and Tween 20 (10 mM Tris-HCL, 150 mM NaCl, 2.5 mM Ca^2+^, 0.1% Tween 20, pH 7.4) for 2 h at RT. The beads were washed 5 times in PBS/0.1% BSA and eluted with 1x LDS buffer for 5 min at 90 °C. Reducing agent was added to the supernatant before loading on a 4–12% bis-tris polyacrylamide gel and running SDS gel electrophoresis using MOPS running buffer. Afterwards, proteins were blotted onto nitrocellulose membranes, subsequently blocked with skim milk. The primary detection antibodies were incubated ON with the blot at 4 °C and the secondary for 1 hr at RT. MBL detection:Hyb 131-01/Rabbit anti-mouse HRP (P0260). C1q detection:A0136/goat anti-rabbit HRP (P0448). The blot was thoroughly washed and developed using SuperSignal West Femto Chemiluminescent Substrate and analyzed on a LI-COR Odyssey XF instrument.

### Immunoprecipitation and western blot detection of C1q/MASP complexes

4 µg/ml F(ab’)2 fragments of anti-MASP-3 mab 7D8, MASP-1/-3/MAP-1 mab 8B3 or F(ab’)2 fragments generated from an irrelevant mouse IgG1 mAb clone MOPC-31C (BD Biosciences, 557273) were coated on maxisorp plates at 4 °C ON and afterwards washed and blocked for 1 hr in TBS/T. 30% serum was added for 2 h at RT, and the plates were washed thoroughly afterwards. Any bound proteins were eluted by adding a 1:1 mix of 10 mM TBS and LDS buffer (Life Technologies) to the first well and then transferring from well to well. The final eluate of 2 rows was used for SDS gel electrophoresis under reducing conditions on a 4–12% bis-tris polyacrylamide gel (Life Technologies, NP0321BOX), which was blotted onto nitrocellulose membranes (Invitrogen, LC2001). Membranes were blocked shortly in skim milk and incubated with the biotin-tagged polyclonal anti-C1q Ab A0136 and HRP-streptavidin for 1h at RT. The blots were developed using SuperSignal West Femto Chemiluminescent Substrate (Thermo Fisher Scientific) and analyzed on a LI-COR Odyssey XF instrument.

### Western blot of the C1 complex

C1 complexes (CompTech, A098) were run in SDS gel electrophoresis on a 4–12% bis-tris polyacrylamide gel under reducing and non-reducing conditions and blotted onto nitrocellulose membranes. Membranes were blocked in skim milk and incubated with anti-MASP-3 mAb 38:12-3 (Hycult Biotech, HM2216) for 1.5h at RT and HRP-conjugated rabbit anti-rat (Agilent, P0450) for 1h at RT. The blots were developed using SuperSignal West Femto Chemiluminescent Substrate and analyzed on a LI-COR Odyssey XF instrument.

### ELISA measurements of C1q/MASP-3, C1q and MASP-3 in a healthy cohort

NHS from a cohort of 99 healthy donors was applied in three different assays to measure C1q/MASP-3, MASP-3 and C1q. To measure C1q/MASP-3 complexes, we used the sandwich ELISA described in ‘*ELISA measurements of serum complexeś* using the F(ab’)2 of an anti-MASP-3 mAb 7D8 as coating Ab. Serum, in a 6.25% or 25% concentration was used as analyte. A serum pool was used as a calibrator to generate arbitrary units to quantify complex levels. MASP-3 levels were measured using anti-MASP-3 mAb 7D8 for coating and a biotin-tagged anti-MASP-1/-3/MAP-1 mAb 8B3 for detection and 5% serum was applied as analyte. rMASP-3 (in-house) was used as calibrator. C1q levels were measured using anti-C1q pAb A0136 for coating and a biotin-tagged anti-C1q pAb A0136 as detection and serum was applied in 0,005%. Purified C1q was used as a calibrator. PBS/0,05% tween 20 was used for washing and dilutions in all assays.

### Pro-FD cleavage by C1q/MASP-3 and MBL/MASP-3

A total of 10 µg/ml rHA were coated on an ELISA plate at 4 °C ON and afterwards incubated with rabbit anti-HA antibody (1:2000 dilution) (Dako, A0001) 2h at RT. Then 1 µg/ml purified C1q was incubated on the plate for 2 h at RT and afterwards 1 µg/ml active MASP-3, zymogen MASP-3 or buffer were incubated at 4 °C ON. Next, 100 µl of pro-FD supernatant from the Flp-In CHO cells were added for 0, 2, 6 and 24 h at 37 °C while shaking. After incubation, 6 µl of the supernatant from each well were run on an SDS PAGE 4–12% bis-tris gel using MES buffer and blotted to a nitrocellulose membrane afterward. Detection of FD was performed with 0.1 µg/ml of an in-house mouse anti-factor D mAb specifically recognizing mature FD and not pro-FD (clone Act 17) and a mAb recognizing all FD (clone F1-11). Rabbit anti-mouse HRP (Dako, P0260) was used as a secondary antibody. Barbital/tween buffer was used for dilution and wash buffer in both assays.

### Complement activation via C1q/rMASP-2 complexes

A total of 10 µg/ml of rHA were coated for 5 h at RT and then incubated with anti-HA antibody 1:2000 at 4 °C ON. A serial dilution of purified C1q was added starting in 1 µg/ml. Cell culture supernatant containing rMASP-2 was added in a dilution of 1:10 for 1h at RT. Afterwards, 4 µg/ml purified C4 (CompTech, A105) or 1% C1q depleted were incubated on the plate for 30 min at RT and C4b deposition was measured as described above. Background signal (no C1q) was subtracted from the assay using purified C4. All dilutions and washing were performed in barbital/tween buffer.

### Zymogen MASP-3 inhibition of classical pathway

10 µg/ml ml rHA were coated for 5 h at RT and afterwards incubated with anti-HA antibody 1:2000 at 4 °C ON. 2 µg/ml purified C1q was added for 2 h at RT and afterwards rMASP-3 was added in a serial dilution for 1 hr at RT. Next, a serial dilution of C1q depleted serum (CompTech, A300) was incubated on the plate for 30 min at 37 °C. C4b deposition was measured using a biotin-tagged anti-C4c polyclonal Ab (Agilent, Q0369) followed by HRP-streptavidin and TMB One. All dilutions and washing were performed in barbital/tween buffer.

### Ethics

Human whole blood was obtained from 99 anonymous donors via the Blood Bank at Rigshospitalet in Copenhagen. Informed written consent was not obtained and required because of an approved exemption from the Regional Ethical Committee in the Capital Region of Denmark and the National Ethical Committee to use blood from anonymous Danish donors for research. All procedures involving the handling of human samples are in accordance with the principles described in the Declaration of Helsinki and ethically approved by the Regional Ethical Committee of the Capital Region of Denmark (H-20028627).

The experimental animal procedures to develop monoclonal antibodies in mice have been approved by the Danish Animal Experiments Inspectorate with the approval ID:2019-15-0201-00090.

## Figure legends

**Suppl. fig. 1. C1q binding to coated F(ab’)2 fragments.** Binding of C1q to coated F(ab’)2 fragments in an ELISA plate was tested to validate the measurements of C1q/MASP-3 complexes. C1q binds to some extent to coated F(ab’)2 fragments (A-C) and directly to the plate (D). However, C1q binding is not affected by EDTA (A-D), confirming the detection of Ca^2+^ dependent complexes in fig. 3.

**Suppl. fig. 2. Anti-FD mAb specifically recognizing mature FD.** Recombinant pro-FD was produced in Flp-In^TM^-CHO cells with or without a protease inhibitor cocktail (PIC) in the culture media creating pro-FD and mature FD, repecetively. A mAb specific against mature FD (Act 17) was generated in mice immunized with a mature FD-peptide and contra-screened against a pro-FD-peptide. The western blot shows detection of rpro-FD and mature FD and the loading control containing mature purified FD (Comptech) using the anti-FD mAb against mature FD (Act 17) (top panel) and a mAb against all FD (F1-11) (bottom panel).

## References

1. Bohlson, Suzanne S., Peter Garred, Claudia Kemper, and Andrea J. Tenner. 2019. “Complement Nomenclature-Deconvoluted.” Frontiers in Immunology 10:1–6.

2. Chen, C B, and R Wallis. 2001. “Stoichiometry of Complexes between Mannose-Binding Protein and Its Associated Serine Proteases. Defining Functional Units for Complement Activation.” The Journal of biological chemistry 276(28):25894–902.

3. Cortesio, Christa L., and Weiping Jiang. 2006. “Mannan-Binding Lectin-Associated Serine Protease 3 Cleaves Synthetic Peptides and Insulin-like Growth Factor-Binding Protein 5.” Archives of Biochemistry and Biophysics 449(1–2):164–70.

4. Dobó, József et al. 2016. “MASP-3 Is the Exclusive pro-Factor D Activator in Resting Blood:The Lectin and the Alternative Complement Pathways Are Fundamentally Linked.” Scientific Reports 6:31877.

5. Garred, Peter et al. 2016. “A Journey through the Lectin Pathway of Complement-MBL and Beyond.” Immunological Reviews 274(1):74–97. http://doi.wiley.com/10.1111/imr.12468.

6. Hayashi, Manabu et al. 2019. “Cutting Edge:Role of MASP-3 in the Physiological Activation of Factor D of the Alternative Complement Pathway.” The Journal of Immunology 203(6):1411–16.

7. Kishore, Uday, and Kenneth B.M. Reid. 2000. “C1q:Structure, Function, and Receptors.” Immunopharmacology 49(1–2):159–70.

8. Kusakari, Kohei et al. 2022. “The Complex Formation of MASP-3 with Pattern Recognition Molecules of the Lectin Complement Pathway Retains MASP-3 in the Circulation.” Frontiers in Immunology 13.

9. Larsen, Flemming et al. 2004. “Disease-Associated Mutations in Human Mannose-Binding Lectin Compromise Oligomerization and Activity of the Final Protein.” The Journal of biological chemistry 279(20):21302–11.

10. Lu, J H et al. 1990. “Binding of the Pentamer/Hexamer Forms of Mannan-Binding Protein to Zymosan Activates the Proenzyme C1r2C1s2 Complex, of the Classical Pathway of Complement, without Involvement of C1q.” The Journal of Immunology 144(6):2287–94.

11. Matsushita, M et al. 2000. “Proteolytic Activities of Two Types of Mannose-Binding Lectin-Associated Serine Protease.” Journal of immunology 165(5):2637–42.

12. Ohta, M., M. Okada, I. Yamashina, and T. Kawasaki. 1990. “The Mechanism of Carbohydrate-Mediated Complement Activation by the Serum Mannan-Binding Protein.” Journal of Biological Chemistry 265(4):1980–84.

13. Oroszlán, Gábor et al. 2015. “MASP-1 and MASP-2 Do Not Activate Pro-Factor D in Resting Human Blood, Whereas MASP-3 Is a Potential Activator:Kinetic Analysis Involving Specific MASP-1 and MASP-2 Inhibitors.” Journal of immunology (Baltimore, Md.:1950) 196(2):857–65.

14. Peter, Maximilian et al. 2022. “Lectin Pathway Enzyme MASP-2 and Downstream Complement Activation in COVID-19.” Journal of Innate Immunity.

15. Phillips, Anna E. et al. 2009. “Analogous Interactions in Initiating Complexes of the Classical and Lectin Pathways of Complement.” The Journal of Immunology 182(12):7708–17.

16. Ricklin, Daniel, George Hajishengallis, Kun Yang, and John D Lambris. 2010. “Complement:A Key System for Immune Surveillance and Homeostasis.” Nature immunology 11(9):785–97.

17. Rosbjerg, Anne, Lea Munthe-Fog, Peter Garred, and Mikkel-Ole Skjoedt. 2014. “Heterocomplex Formation between MBL/Ficolin/CL-11-Associated Serine Protease-1 and-3 and MBL/Ficolin/CL-11-Associated Protein-1.” Journal of immunology 192(9):4352–60.

18. Skjoedt, Mikkel-Ole, Tina Hummelshoj, et al. 2010. “A Novel Mannose-Binding Lectin/Ficolin-Associated Protein Is Highly Expressed in Heart and Skeletal Muscle Tissues and Inhibits Complement Activation.” The Journal of biological chemistry 285(11):8234–43.

19. Skjoedt, Mikkel-Ole, Yaseelan Palarasah, et al. 2010. “MBL-Associated Serine Protease-3 Circulates in High Serum Concentrations Predominantly in Complex with Ficolin-3 and Regulates Ficolin-3 Mediated Complement Activation.” Immunobiology 215(11):921–31.

20. Teillet, Florence et al. 2005. “The Two Major Oligomeric Forms of Human Mannan-Binding Lectin:Chemical Characterization, Carbohydrate-Binding Properties, and Interaction with MBL-Associated Serine Proteases.” The Journal of Immunology 174:2870–77.

21. Thiel, S et al. 2000. “Interaction of C1q and Mannan-Binding Lectin (MBL) with C1r, C1s, MBL-Associated Serine Proteases 1 and 2, and the MBL-Associated Protein MAp19.” Journal of immunology (Baltimore, Md.:1950) 165(2):878–87.

22. Thielens, Nicole M., F. Tedesco, Suzanne S. Bohlson, and Christine Garboriaud. 2017. “C1q:A Fresh Look upon an Old Molecule.” Molecular Immunology (89):73–83.

