## Supplementary figures and images for "C1q/MASP complexes – hybrid complexes of classical and lectin pathway proteins are found in the circulation"

### suppl. fig. 1

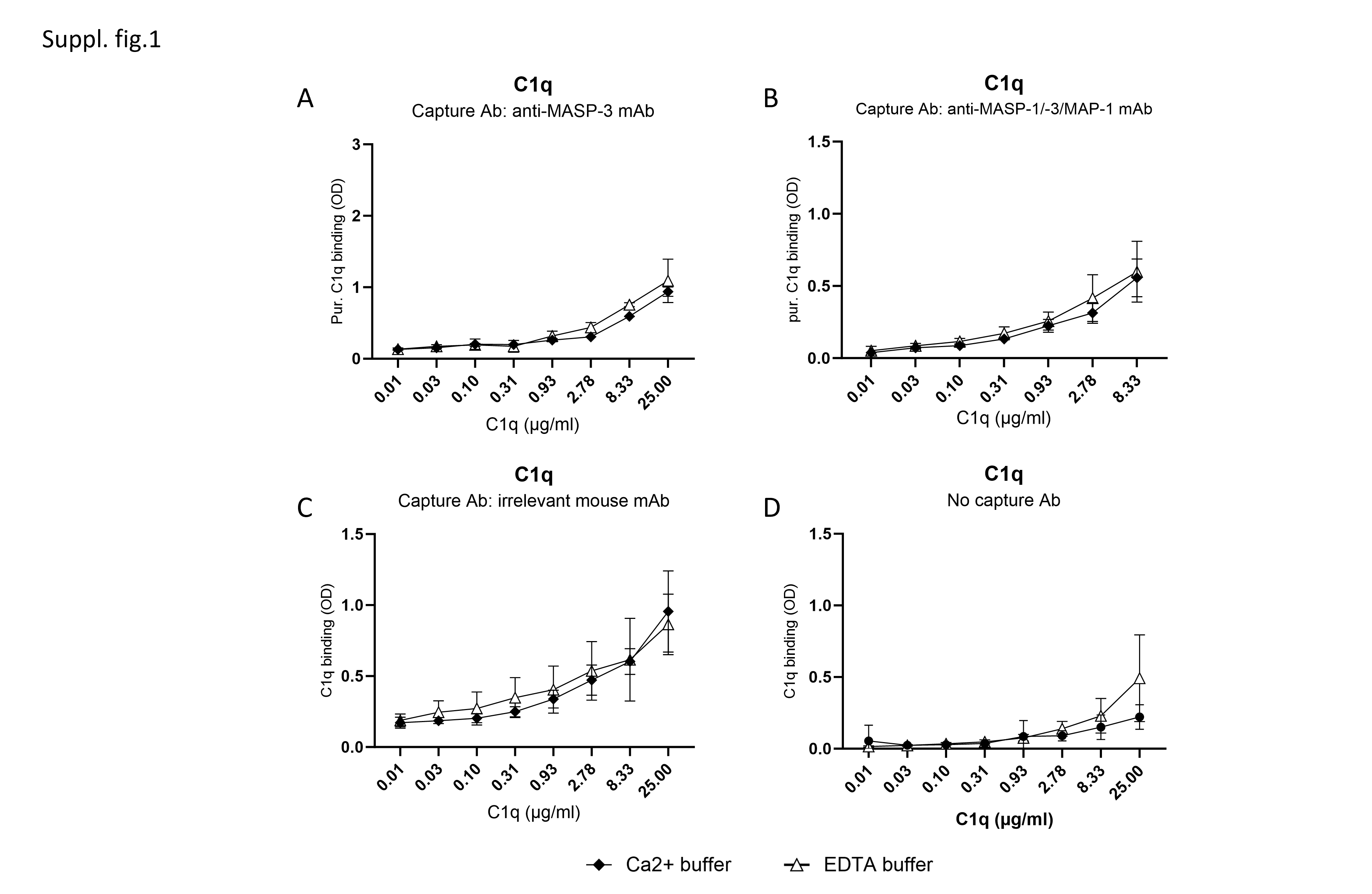

### suppl. fig. 2

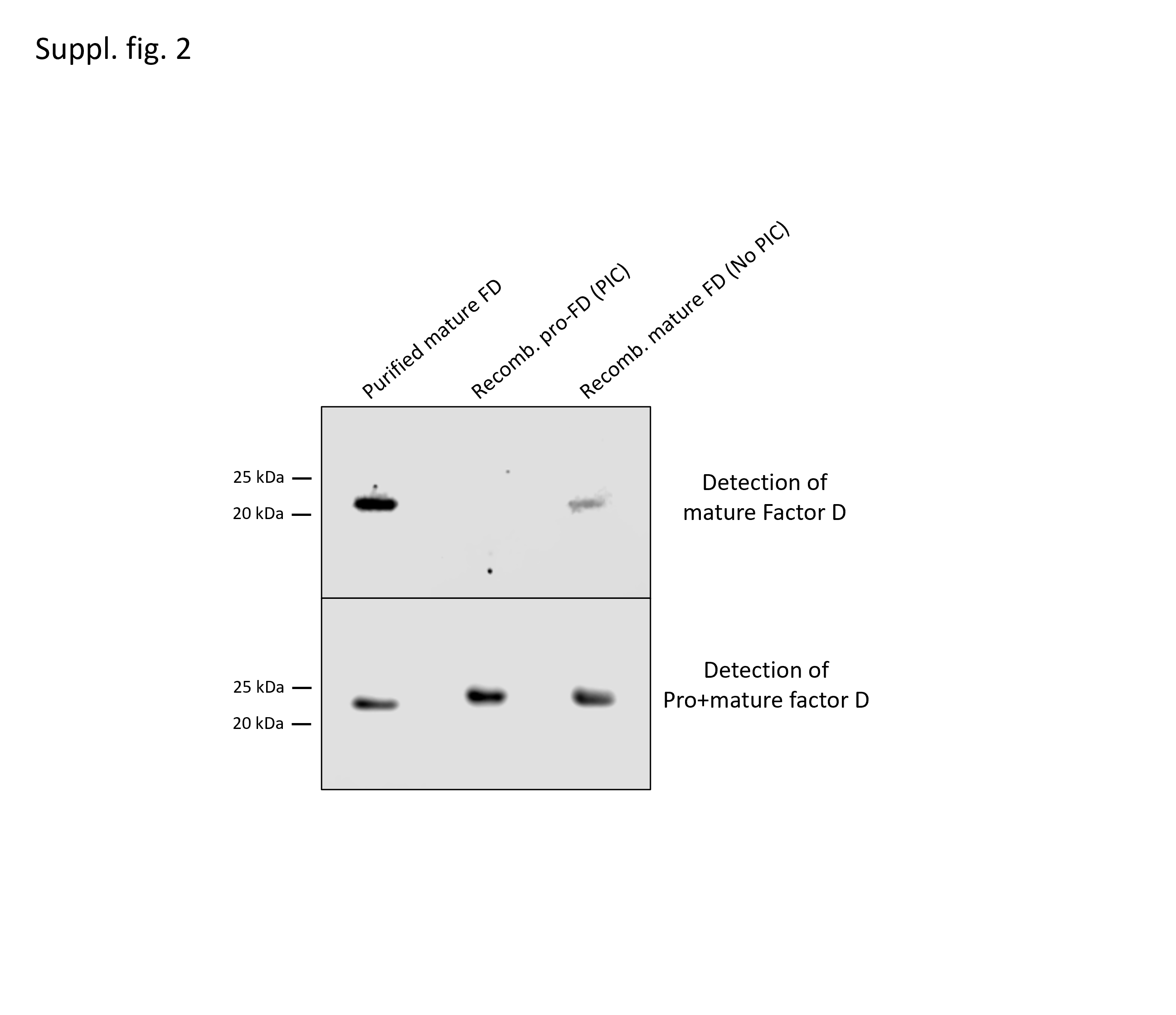
